# Vascular-Derived SPARC and SerpinE1 Regulate Interneuron Tangential Migration and Accelerate Functional Maturation of Human Stem Cell-Derived Interneurons

**DOI:** 10.1101/2020.02.07.939298

**Authors:** Matthieu Genestine, Daisy Ambriz, Gregg W. Crabtree, Anna Molotkova, Michael Quintero, Angeliki Mela, Saptarshi Biswas, Peter Canoll, Gunnar Hargus, Dritan Agalliu, Joseph A. Gogos, Edmund Au

## Abstract

Cortical interneurons establish inhibitory microcircuits throughout the neocortex and their dysfunction has been implicated in epilepsy and neuropsychiatric diseases. Developmentally, interneurons migrate from a distal progenitor domain in order to populate the neocortex – a process that occurs at a slower rate in humans than in mice. In this study, we sought to identify factors that regulate the rate of interneuron maturation across the two species. Using embryonic mouse development as a model system, we found that the process of initiating interneuron migration is regulated by blood vessels of the medial ganglionic eminence (MGE), an interneuron progenitor domain. We identified two endothelial cell-derived paracrine factors, SPARC and SerpinE1, that enhance interneuron migration in mouse MGE explants and organotypic cultures. Moreover, pre-treatment of human stem cell-derived interneurons (hSC-interneurons) with SPARC and SerpinE1 prior to transplantation into neonatal mouse cortex enhanced their migration and morphological elaboration in the host cortex. Further, SPARC and SerpinE1-treated hSC-interneurons also exhibited more mature electrophysiological characteristics compared to controls. Overall, our studies suggest a critical role for CNS vasculature in regulating interneuron developmental maturation in both mice and humans.

## Introduction

Cortical interneurons are inhibitory, locally projecting cells that form a distributed network of repetitive microcircuits throughout the cortex (Kepecs and Fishell, 2014; Tremblay et al., 2016). To establish this distributed network, interneurons migrate from the ventral subpallium to arealize throughout the cortex in order to form connections with layered pyramidal neurons and other interneurons. The timescale for interneuron migration varies widely across species; The process lasts days in mice (Lavdas et al., 1999; Marin and Rubenstein, 2001) and several months in humans (Arshad et al., 2016; Hansen et al., 2013; Ma et al., 2013). Further, this species difference extends to stem cell derived interneurons where mouse ES-derived interneurons develop rapidly (Au et al., 2013; Maroof et al., 2010; McKenzie et al., 2019), while hSC-interneuron development is protracted (Nicholas et al., 2013; Shao et al., 2019). Indeed, this limitation has restricted the scope of analysis on hSC-interneurons, hampering detailed functional studies.

In this study, we sought to identify factors that regulate the timing of interneuron migration across species. Using embryonic mouse development as a model system, we found that interneuron migration coincides with vascularization of the medial ganglionic eminence (MGE). In fact, by genetically manipulating the degree of MGE vascularization *in vivo*, we can alter the degree of interneuron migration into the cortex. Using an in vitro approach, we identified paracrine factors, SPARC and SerpinE1 produced by endothelial cells that enhance interneuron migration in mouse MGE explants and organotypic cultures. Given the slow developmental rate of hSC-interneurons, we tested whether SPARC and SerpinE1 treatment similarly induced migration in human cells. We found that treated hSC-interneurons xenografted into neonatal mouse cortex exhibited enhanced migration. Further, transplanted hSC-interneurons also showed greater morphological complexity, and more mature electrophysiological characteristics compared to controls. These data suggest that interneurons in mice and humans share a common vascular-based mechanism that regulates their developmental timing. Notably, by characterizing this process in mice, we were able to reverse-engineer our findings to accelerate functional maturation in human interneurons.

## Results

Cortical interneurons are generated from a distal source (primarily the medial gangionic eminence, MGE) and undergo long distance migration to their final destination in the neocortex. In the embryonic mouse, this developmental process is rapid: interneurons are generated starting approximately e11 and robustly migrate into the cortex by e15 (Figure 1A) (Lavdas et al., 1999; Marin and Rubenstein, 2001). As a result, it is tempting to assume that newly born interneurons automatically transition to a migratory state. As a counter example, however, in human fetal development interneuron migration into the cortex is highly protracted. Previous studies have found that postmitotic interneurons slowly transition to a migratory state to populate the neocortex over the course of many weeks (Arshad et al., 2016; Hansen et al., 2013; Ma et al., 2013). To confirm this, we examined interneurons in fetal cortical sections by immunohistochemistry for Dlx2 (an interneuron marker). Consistent with prior reports, we observed an increase in the density of Dlx2^+^ interneurons in the cortex over time (15 post conception weeks (pcw) to 22 pcw) (Figure 1B, Figure S1A,B). We hypothesized that the discrepancy between human and mouse interneuron development may be due to an external cue delivered to the MGE to promote interneuron migration. We therefore examined the developing mouse MGE (from e10.5 to e15.5) by histology and observed a striking increase in vascularization of the MGE and the underlying mantle region during the time period when mouse interneuron migration initiates (Figure 1C, D) (Daneman et al., 2009; Paredes et al., 2018). Similarly, we found that vascularization of human fetal MGE slowly increased with developmental age, with robust vascularization not occurring until after 20 pcw (Figure 1E, Figure S1C,D).

**Figure 1.**
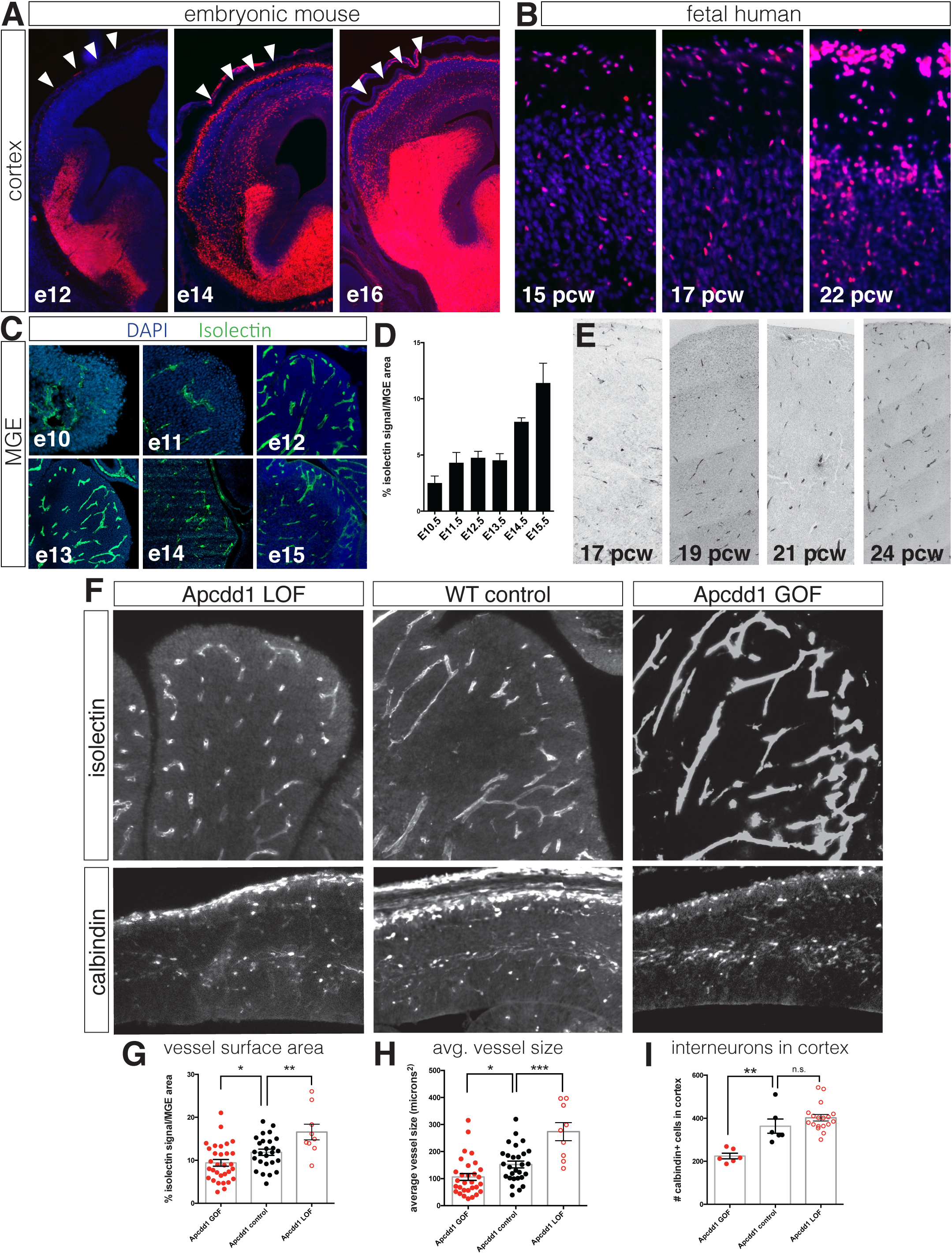
Interneuron migration progresses over development and is regulated by MGE vascularization. (A) Coronal sections of Dlx6a^Cre^; Ai9 embryonic mouse telencephalon over developmental time. tdTomato+ interneurons progressively migrate into the cortex between e12 and e16. Cortex indicated by arrowheads. (B) Dlx2 immunohistochemistry (red) in human fetal cortex show weeks-long progression of cortical interneurons migrating into cortex. (C) Representative images of isolectin labeling (green). (D) Quantification of isolectin+ blood vessel staining as a percentage of MGE area for embryos ages e10.5 to e15.5. (E) CD31 immunohistochemistry in human MGE from various fetal ages. (F) Top row, coronal sections of e14.5 MGE. Blood vessels labeled with isolectin in wildtype control, *Apcdd1* loss-of-function and *Apcdd1* gain-of-function mutants. Bottom row, coronal sections of e14.5 embryonic brain labeled with calbindin to show migratory interneurons in isolectin in wildtype control, *Apcdd1* loss-of-function and *Apcdd1* gain-of-function mutants. (G, H) Quantification of isolectin+ MGE vascularization; (G) isolectin+ vessel labeling as a percentage of MGE surface area; (H) average vessel size in square microns. (I) Quantification of total interneurons migrating into cortex/20um section. Paired t-test, * p < 0.05; **p < 0.01; *** p < 0.001.

To determine whether there is a causal relationship between MGE vascularization and interneuron migration, we examined *Apcdd1* mutant mice (Mazzoni et al., 2017). Apcdd1 a negative regulator of Wnt/β-catenin signaling, that is critical for CNS vascular development and blood-brain barrier maturation (Shimomura et al., 2010). We examined MGE vessel density and average vessel size in *Apcdd1* GOF and LOF mutants at e14.5 and found that, similar to the retina and cerebellum (Mazzoni et al., 2017), vascularization in the MGE is decreased in *Apcdd1* GOF and increased in *Apcdd1* LOF mice versus wildtype controls (Figure 1F-H). Importantly, fewer interneurons migrated into the cortex in Apcdd1 GOF mutants, in which vascularization was less extensive compared to wild type controls. Conversely, a trend towards more Calbindin+ migrating interneurons were present in *Apcdd1* LOF mutants (p=0.236, paired t-test) in which vessel size and density was greater (Figure 1I). These results suggest that the density of endothelial cells within neural tissue is a critical regulator interneuron migration.

In order to identify a mechanism by which this occurs, we used an MGE explant culture approach from mouse embryos to examine the number of migrating neurons at various developmental stages (Figure 2A, Figure S2A). Given that the MGE is progressively vascularized from e10 to e15, we reasoned that MGE explants may exhibit a developmental age-dependent capacity to migrate from the explant. We further reasoned that in early MGE explants that were not extensively vascularized that mouse interneurons would be similarly immobile like human interneurons. To normalize for differences in MGE size at different ages, whole MGE tissue was dissected and sectioned at 250 microns. Then, the number of DAPI+ migratory cells was normalized to the surface area of the MGE explant. Using this approach, we found a strong linear correlation between the number of migratory cells and MGE explant surface area (r^2^=0.6829) (Figure S2B). Further, we found that nearly all migratory DAPI+ cells were interneurons using the Dlx6a^Cre^ driver line crossed to Ai9 in order to fate map the migratory lineage (Monory et al., 2006) (Figure S2C,D). Classic studies have shown that MGE explants at e14.5 and e15.5 exhibit robust interneuron migration within hours of initial plating (Bellion et al., 2005; Polleux et al., 2002; Wichterle et al., 1999). Consistently, we found that interneurons later time point MGE explants exhibited robust migration, whereas explants from earlier time points (e10.5 to e12.5) had a limited capacity to migrate (Figure 2B, Figure S2C). One possibility is that interneurons possess an intrinsic timer such that, given sufficient time in culture, early timepoint explants would migrate to the same extent as older MGE explants. However, even when cultured for up to a week, early explants did not significantly migrate more after the first 48 hours (data not shown).

**Figure 2.**
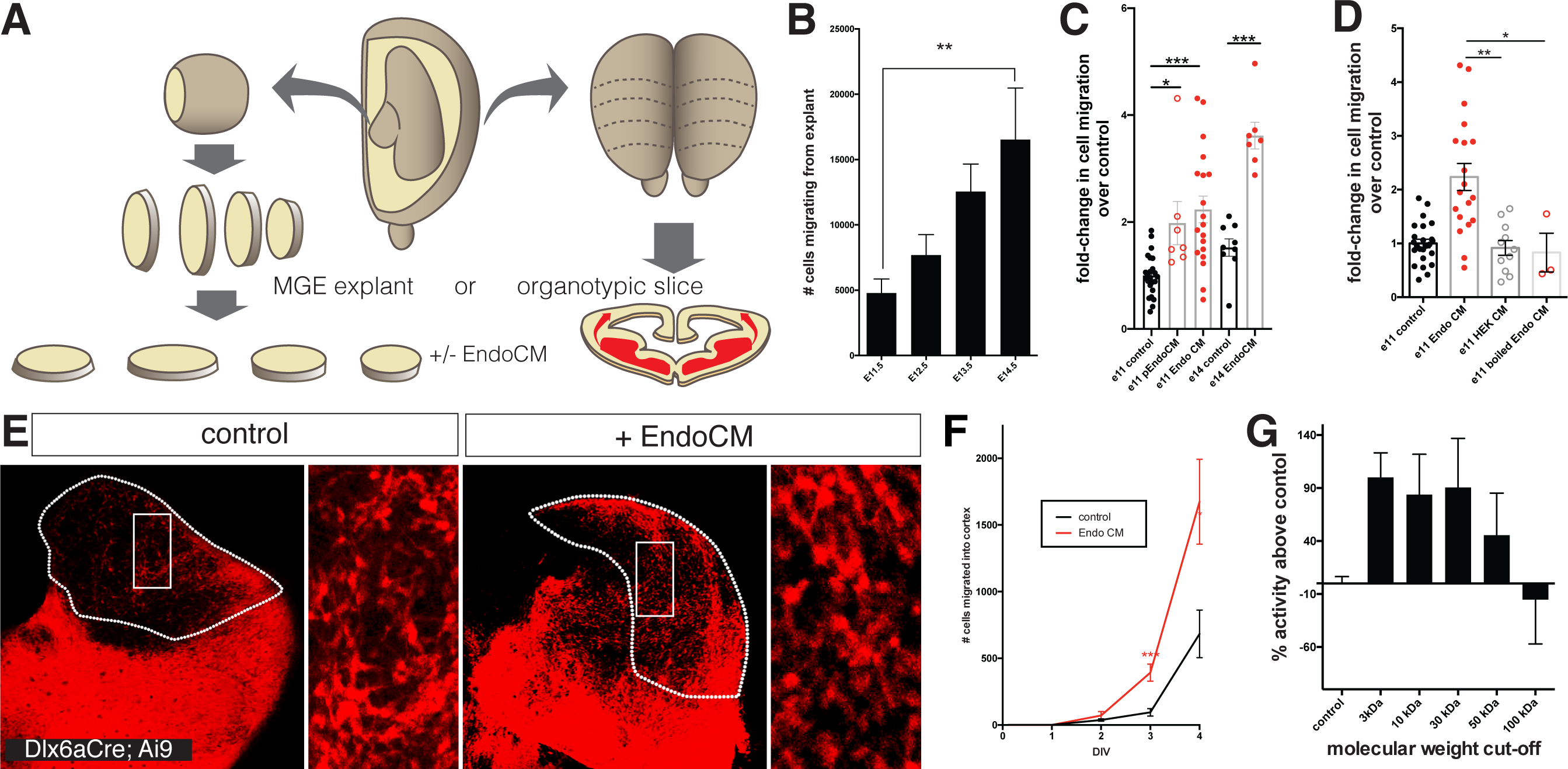
Endothelial cell conditioned medium increases interneuron migration in MGE explants and organotypic slice cultures. (A) Schematic representation of MGE explant and organotypic slice preparation to assess interneuron migration *in vitro*. (B) Number of interneurons (normalized to MGE surface area) migrating from MGE explants from embryos ages e10.5 to e15.5. (C) Number of interneurons (normalized to MGE surface area) migrating from e11.5 and e14.5 MGE explants with or without primary culture endothelial cell conditioned medium (p-EndoCM) or immortalized endothelial cell line conditioned medium (EndoCM). (D) Number of interneurons (normalized to MGE S.A.) migrating from e11.5 MGE explants treated with control, EndoCM, HEK293 conditioned medium (HEKCM) or boiled EndoCM. (E) Representative images of DIV4 organotypic slice cultures without (control) or with EndoCM added. Right is higher magnification of boxed region on left. (F) Number of Dlx6a^Cre^; Ai9 tdTomato+ interneurons migrating into cortex over time in coronally-section organotypic slice cultures (DIV 0 to 4) with or without EndoCM treatment. (G) Size fractionation of EndoCM medium assayed for normalized interneuron migration from e11.5 MGE explants. Paired t-test, * p < 0.05; **p < 0.01; *** p < 0.001.

Given the diffuse distribution of blood vessels in the MGE, the most likely mechanism is that endothelial cells produce a paracrine signal to induce interneuron migration. To test this directly, we prepared conditioned medium from primary cultures of embryonic brain ECs (p-EndoCM) and added it to e11.5 MGE explants. Consistent with our hypothesis, addition of p-EndoCM resulted in a robust increase in interneuron migration (Figure 2C). We found, however, that primary embryonic brain EC cultures introduced unwanted variability, and for subsequent experiments, we obtained Endo-CM from an immortalized human EC line, HBEC5i (Wassmer et al., 2006). To further reduce inter-experimental variability, interneuron migration counts were normalized to within-experiment negative (untreated) controls. Hence, subsequent data is presented as fold-change in interneuron migration over controls. Similar to p-Endo CM, conditioned medium from HBEC-5i (Endo CM) also robustly increased e11.5 MGE explant migration (Figure 2C). Interestingly, e14.5 MGE migration was also significantly increased by EndoCM (Figure 2C). We next tested the effect of EndoCM on interneuron migration in an organotypic slice culture. Here, we used Dlx6a^Cre^; Ai9+ e11.5 embryos in order to visualize interneurons as they migrate within a coronal slice of telencephalon from the MGE into the cortex (Figure 2A,E). As with MGE explants, EndoCM also significantly increased the rate and overall number of interneurons that migrated into the cortex (Figure 2F). As a negative control, we tested the biological activity of conditioned medium from HEK 293 cells (HEK CM). HEK CM did not increase MGE explant migration at either age (Figure 2D). Moreover, we found that pre-boiling EndoCM eliminated its biological activity, suggesting a protein source as a regulator of interneuron migration (Figure 2D). Finally, we size-fractionated EndoCM and found that biological activity was strongly reduced between 30 kDa and 100 kDa (Figure 2G).

In order to identify candidate proteins, we performed bulk RNA sequence analysis on HBEC5i and HEK cells. We screened the dataset for genes with the greatest differential expression, enriched in HBEC5i that produced proteins between 30 and 100 kDa, which were also secreted (GO term: extracellular space) (Figure S3). After further curation to eliminate membrane-tethered molecules, we obtained a short list of 24 candidates (Table 1). We functionally tested a number of candidates, including VEGF-A (Barber et al., 2018) and Follistatin; however, two proteins, SPARC and Serpin E1 exhibited the most robust biological activity in their ability to increase e11 MGE migration (Figure 3).

**Figure 3.**
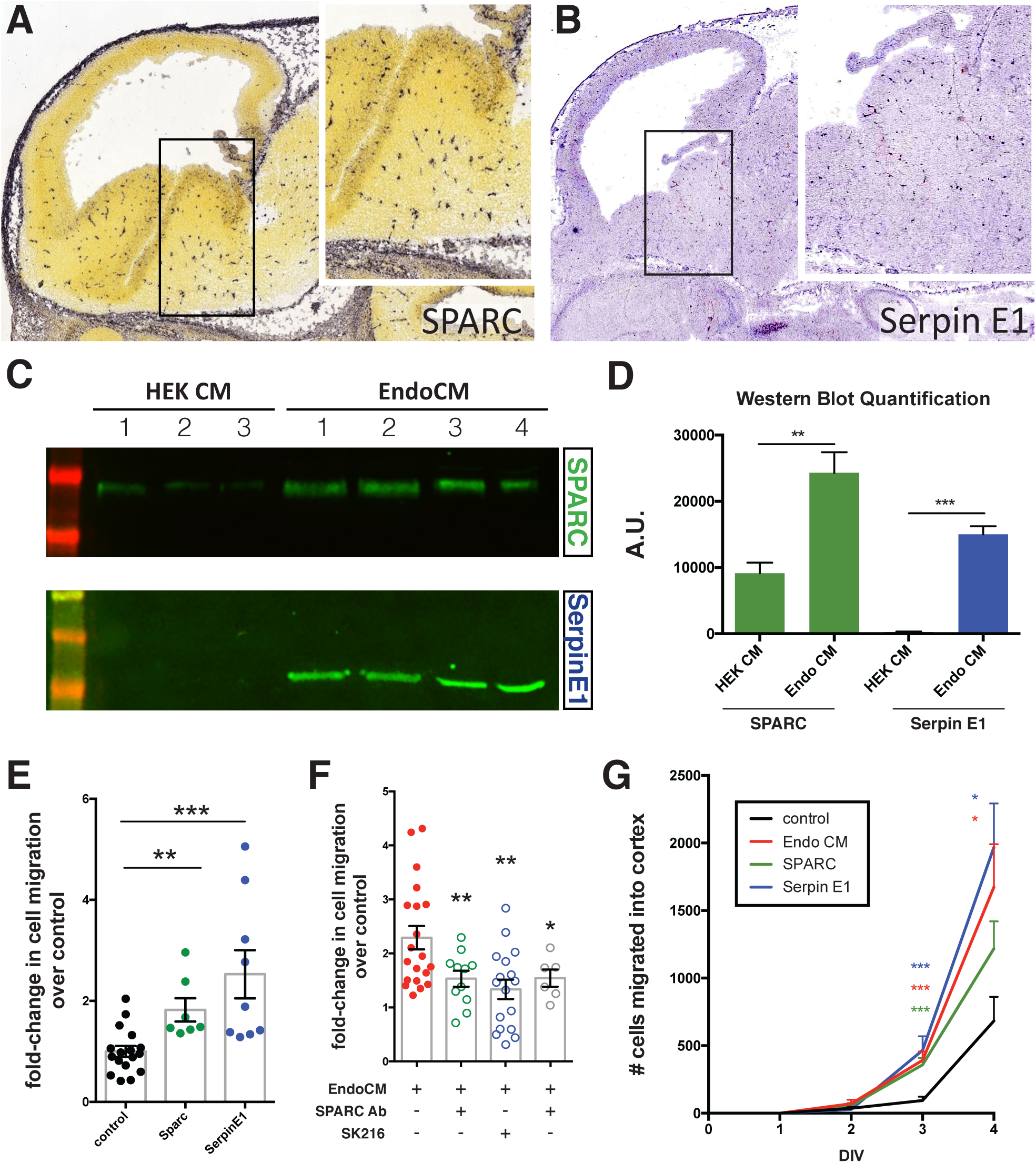
Endothelially-derived factors, SPARC and SerpinE1 increase interneuron migration and account for most of the biological activity of EndoCM. (A, B) Parasaggital sections of e13.5 embryonic brain showing in situ hybridization signal for (A) SPARC* and (B) Serpin E1^#^. Insets for (a) and (b) are higher magnification or boxed regions showing MGE expression. (C) Western blots of HEK CM (3 replicates) and EndoCM (4 replicates) for SPARC and SerpinE1. (D) Quantification of western band intensity for SPARC (green) and SerpinE1 (blue) in HEK CM and EndoCM. (E) Quantification of normalized interneuron migration from MGE explants treated with SPARC (filled green), Serpin E1 (filled blue) compared with control (filled black). (F) Quantification of normalized interneuron migration from MGE explants treated with EndoCM (filled red), EndoCM and SPARC function-blocking antibody (unfilled green), EndoCM and SerpinE1 small molecule inhibitor (unfilled blue) and EndoCM with combination of SPARC function-blocking antibody and Serpin E1 small molecule inhibitor (unfilled gray). (G) Number of Dlx6a^Cre^; Ai9 tdTomato+ interneurons migrating into cortex over time in coronally-section organotypic slice cultures (DIV 0 to 4) with or without EndoCM, SPARC or SerpinE1 treatment. Paired t-test, * p < 0.05; **p < 0.01; *** p < 0.001. * from Allen Brain Atlas (http://developingmouse.brain-map.org/); # from Gene Paint (http://gp3.mpg.de).

**Table 1.**
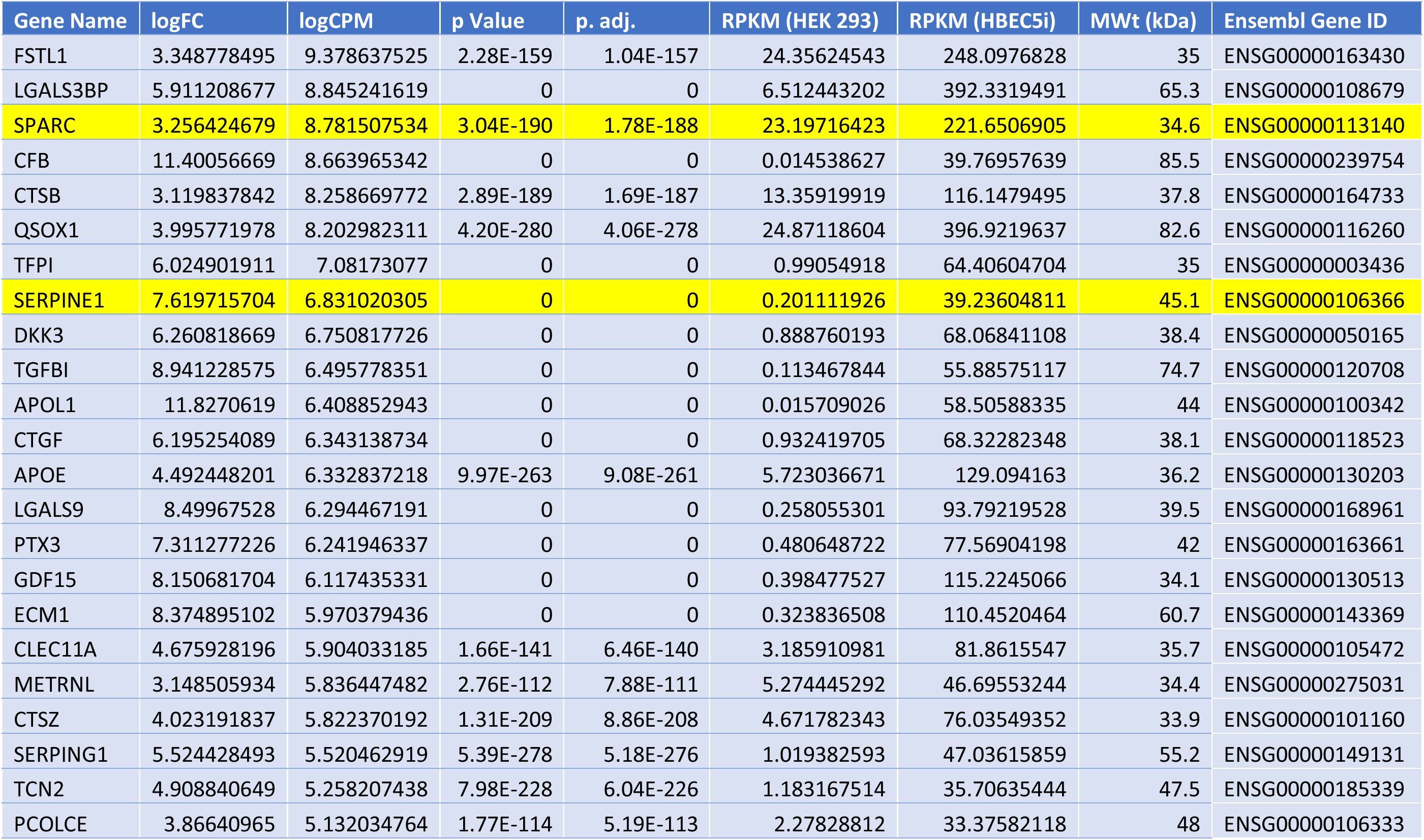
Short List of Candidate Factors

| Gene Name | logFC | logCPM | p Value | p. adj. | RPKM (HEK 293) | RPKM (HBEC5i) | MWt (kDa) | Ensembl Gene ID |
| --- | --- | --- | --- | --- | --- | --- | --- | --- |
| FSTL1 | 3.348778495 | 9.378637525 | 2.28E-159 | 1.04E-157 | 24.35624543 | 248.0976828 | 35 | ENSG00000163430 |
| LGALS3BP | 5.911208677 | 8.845241619 | 0 | 0 | 6.512443202 | 392.3319491 | 65.3 | ENSG00000108679 |
| SPARC | 3.256424679 | 8.781507534 | 3.04E-190 | 1.78E-188 | 23.19716423 | 221.6506905 | 34.6 | ENSG00000113140 |
| CFB | 11.40056669 | 8.663965342 | 0 | 0 | 0.014538627 | 39.76957639 | 85.5 | ENSG00000239754 |
| CTSB | 3.119837842 | 8.258669772 | 2.89E-189 | 1.69E-187 | 13.35919919 | 116.1479495 | 37.8 | ENSG00000164733 |
| QSOX1 | 3.995771978 | 8.202982311 | 4.20E-280 | 4.06E-278 | 24.87118604 | 396.9219637 | 82.6 | ENSG00000116260 |
| TFPI | 6.024901911 | 7.08173077 | 0 | 0 | 0.99054918 | 64.40604704 | 35 | ENSG00000003436 |
| SERPINE1 | 7.619715704 | 6.831020305 | 0 | 0 | 0.201111926 | 39.23604811 | 45.1 | ENSG00000106366 |
| DKK3 | 6.260818669 | 6.750817726 | 0 | 0 | 0.888760193 | 68.06841108 | 38.4 | ENSG00000050165 |
| TGFBI | 8.941228575 | 6.495778351 | 0 | 0 | 0.113467844 | 55.88575117 | 74.7 | ENSG00000120708 |
| APOL1 | 11.8270619 | 6.408852943 | 0 | 0 | 0.015709026 | 58.50588335 | 44 | ENSG00000100342 |
| CTGF | 6.195254089 | 6.343138734 | 0 | 0 | 0.932419705 | 68.32282348 | 38.1 | ENSG00000118523 |
| APOE | 4.492448201 | 6.332837218 | 9.97E-263 | 9.08E-261 | 5.723036671 | 129.094163 | 36.2 | ENSG00000130203 |
| LGALS9 | 8.49967528 | 6.294467191 | 0 | 0 | 0.258055301 | 93.79219528 | 39.5 | ENSG00000168961 |
| PTX3 | 7.311277226 | 6.241946337 | 0 | 0 | 0.480648722 | 77.56904198 | 42 | ENSG00000163661 |
| GDF15 | 8.150681704 | 6.117435331 | 0 | 0 | 0.398477527 | 115.2245066 | 34.1 | ENSG00000130513 |
| ECM1 | 8.374895102 | 5.970379436 | 0 | 0 | 0.323836508 | 110.4520464 | 60.7 | ENSG00000143369 |
| CLEC11A | 4.675928196 | 5.904033185 | 1.66E-141 | 6.46E-140 | 3.185910981 | 81.8615547 | 35.7 | ENSG00000105472 |
| METRNL | 3.148505934 | 5.836447482 | 2.76E-112 | 7.88E-111 | 5.274445292 | 46.69553244 | 34.4 | ENSG00000275031 |
| CTSZ | 4.023191837 | 5.822370192 | 1.31E-209 | 8.86E-208 | 4.671782343 | 76.03549352 | 33.9 | ENSG00000101160 |
| SERPING1 | 5.524428493 | 5.520462919 | 5.39E-278 | 5.18E-276 | 1.019382593 | 47.03615859 | 55.2 | ENSG00000149131 |
| TCN2 | 4.908840649 | 5.258207438 | 7.98E-228 | 6.04E-226 | 1.183167514 | 35.70635444 | 47.5 | ENSG00000185339 |
| PCOLCE | 3.86640965 | 5.132034764 | 1.77E-114 | 5.19E-113 | 2.27828812 | 33.37582118 | 48 | ENSG00000106333 |

We examined the expression of SPARC and Serpin E1 in e14.5 MGE through publicly-available expression databases (Allen Brain Atlas and Gene Paint), and found that both were specifically expressed in the embryonic CNS vasculature (Figure 3A,B). We also confirmed by western blotting that SPARC and SerpinE1 were present in EndoCM at significantly higher levels compared to HEK293 CM (Figure 3C,D). We then added recombinant SPARC or SerpinE1 to MGE explants and found that cell migration was increased, in particular with SerpinE1 (Figure 3E). We then tested EndoCM in which SPARC and Serpin E1 activity were depleted. We used a function-blocking antibody for SPARC (Sweetwyne et al., 2004), whereas Serpin E1 activity was blocked with a small molecule inhibitor (SK216 (Masuda et al., 2013)). These reagents separately significantly reduced the capacity of Endo CM to increase e11.5 MGE migration (Figure 3F). However, inhibition of both SPARC and SerpinE1 did not reduce interneuron migration further, suggest that the two molecules may act on intersecting pathways. Finally, we tested whether SPARC and Serpin E1 could increase interneuron migration in an organotypic slice over time. We found that both proteins significantly increased the rate and overall number of interneurons that migrated into the cortex over time (Figure 3G). SPARC and SerpinE1 have been shown to inhibit both endothelial cell migration and proliferation (Boyineni et al., 2016; Hasselaar and Sage, 1992; Lane et al., 1994; Menashi et al., 1993).

Previous studies have demonstrated that hSC-interneurons migrate and mature at a slow rate, reminiscent of the protracted time frame of interneuron development in the human fetus (Arshad et al., 2016; Hansen et al., 2013; Ma et al., 2013). Given that SPARC and Serpin E1 elicit interneuron migration in e11.5 MGE explants and organotypic slice cultures, we tested whether these factors might similarly accelerate the developmental timeframe for hSC-interneurons. Using a pan-tdTomato expressing human iPSC line (CAG-tdTomato knocked into *AAVS1* locus), we achieved efficient differentiation to ventral telencephalic identity using established protocols (Bagley et al., 2017) (Figure S4). hSC-interneuron differentiation was confirmed using AAV Dlx5/6-GFP(Dimidschstein et al., 2016) (Figure S4B). At day 35 of differentiation (DIFF 35), ventral telencephalic organoids were treated with either SPARC, Serpin E1 or both for 14 days in order to observe a biological effect *in vitro* (data not shown). At DIFF 49, we tested for hSC-interneuron migration by dissociating either untreated and SPARC and SerpinE1-treated organoids. Here, we further divided the groups: one group continued to be exposed to SPARC and SerpinE1 and the other was left untreated (Figure S5A). We observed a significant increase in migratory distance in the treated group. Importantly, we observed an even greater biological effect when SPARC and Serpin were added to dissociated cells after pre-treatment (Figure S7c). Thus, in subsequent xenograft experiments, SPARC and SerpinE1 was added as a pre-treatment for 14 days and also added to the cells at the time of transplantation.

Previous studies have xenografted hSC-interneurons into a more functionally relevant setting: neonatal mouse cortex. They found that functional maturation rate is prolonged, requiring approximately 7 months (Nicholas et al., 2013; Shao et al., 2019). We then tested the capacity of control and SPARC/SerpinE1-treated ventral organoids to integrate following xenotransplant into immune-compromised (NSG) mouse cortex. We first analyzed transplants 28 days post engraftment. We confirmed that tdTomato+ hSC-interneurons were almost all Dlx2+ (Figure 4A), and also found that Combo-treated hSC-interneurons migrated significantly further than controls (Figure 4B,C). At 56 days post-transplantation, we traced and analyzed the morphologies of hSC-interneurons in 3D and found that they possessed longer processes and branched more extensively (Figure 4D, Figure S6).

**Figure 4.**
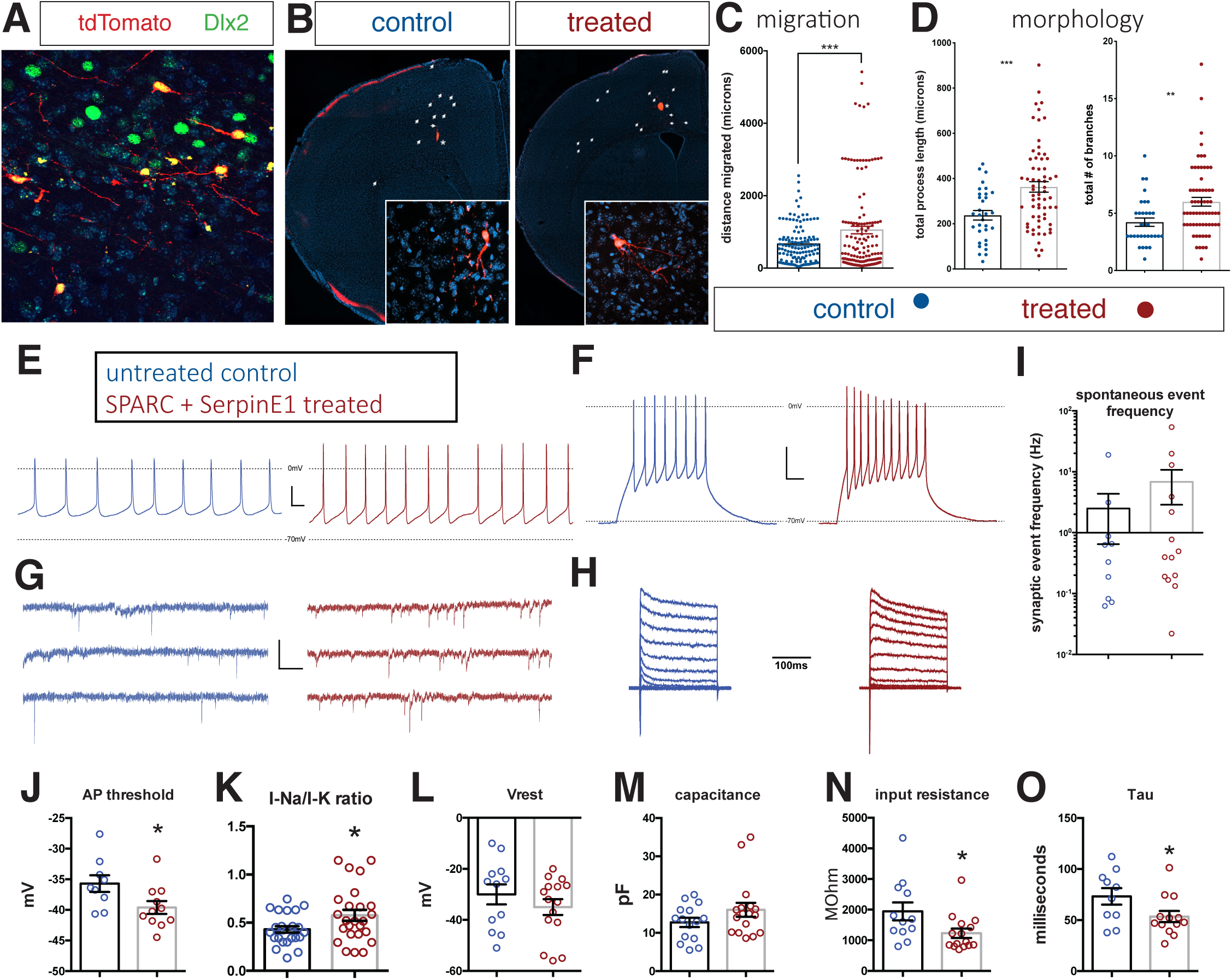
Human stem cell-derived interneurons xenografted into host mouse cortex are more migratory and more morphologically-complex with SPARC and SerpinE1pre-treatment. (A) Immunohistochemistry for Dlx2 (green), Dapi (blue) and tdTomato+ hSC-interneurons (red) near injection site 1-month post-transplant. (B) Representative images of coronal sections of host mouse cortex 1 month post-transplant of control untreated and SPARC/SerpinE1 pre-treated hSC-interneurons (asterisk denotes transplant site). Arrowheads show tdTomato+ hSC-interneurons migrating away from transplant site. (C) Quantification of migratory distance of hSC-interneurons from site of injection 1-month post transplantation. (D) Quantification of hSC-interneuron morphology by average process length (left) and total number of neurite branches (right). (E-O) Whole cell recordings of hSC-interneurons 2-months post-transplant. Untreated control (blue), SPARC and SerpinE1 pre-treated (red). (E) Representative traces of spontaneous AP firing. (F) Evoked firing upon square pulse depolarization. (G) Representative traces of spontaneous miniature EPSCs. (H) Step recordings of sodium current. (I-O) Quantitative measurements of (I) Spontaneous mEPSC frequency (Hz), (J) AP threshold (mV) (p=0.0166), (K) Sodium-to-potassium current ratio (p=0.0338), (L) resting membrane potential (p=0.12), (M) capacitance (p=0.067), (N) input resistance (p=0.0155), (O) Tau (p=0.0239). Paired t-test, * p < 0.05; **p < 0.01; *** p < 0.001. Representative traced morphology of 2-month post-transplant tdTomato+ hSC-interneuron.

Given the improvements in migration and morphology exhibited by treated hSC-interneurons, we tested their functional maturity by employing whole cell recordings from Td-tomato labelled transplanted cells in acute slices of frontal cortex 56 days post-transplant. Consistent with more mature neuronal function, recordings from treatment group neurons showed significantly reduced membrane input resistance, faster membrane time constants, more hyperpolarized action potential thresholds, and increased ratios of voltage gated sodium currents versus voltage gated potassium currents (Figure 4E-O). Additionally, other measures of neuronal maturity we tested appeared to collectively trend towards more mature functional phenotypes in the treatment group of neurons, albeit these were not statistically significant (Figure S7). Specifically, treated cells appear to have larger membrane capacitances (p=0.067), more hyperpolarized resting potentials (p=0.12), larger voltage gated sodium currents (p=0.20), faster AP rise rates (p=0.21), narrower APs (p=0.22), and generated a greater number of APs (p=0.24) (Figure S9). Taken together, these electrophysiological profiles suggest that treated neurons may have an accelerated trajectory towards functional maturity.

## Discussion

CNS vascularization is a tightly regulated process that operates in tandem with neuronal and glial cell development (Paredes et al., 2018). Here, we demonstrate that not only does vascularization of mouse MGE coincide with interneuron migration, but it also plays a regulatory role since genetic manipulation of vascular pruning affects the number of interneurons migrating into the cortex. Further, two EC-derived paracrine factors SPARC and Serpin E1 induce increased interneuron migration as assessed by *in vitro* assays. Having demonstrated a critical crosstalk between the neural and vascular compartments in mouse, we leveraged our findings to test if SPARC and SerpinE1 can accelerate hSC-interneuron maturation. We found that hSC-interneurons treated with SPARC and Serpin E1 also show a more robust migration both *in vitro* and upon xenotransplantation into host mouse cortex. Finally, xenografted hSC-interneurons also exhibit significantly more complex morphologies and some mature electrophysiological properties.

Our findings support previous studies that have linked angiogenesis to pathfinding during interneuron tangential migration (Barber et al., 2018; Li et al., 2018) and MGE mitosis (Tan et al., 2016). Further, radial glia in the MGE undergo a transition during embryonic development where their pial endfeet detach and connect to MGE vasculature (Tan et al., 2016). In light of our data, it is tempting to speculate that this rearrangement allows for a more direct communication between ECs and interneurons, thereby facilitating their tangential migration into the cortex. In humans, interneuron migration during fetal development is a protracted process (Arshad et al., 2016; Hansen et al., 2013; Ma et al., 2013). We found that hSC-interneurons exhibit greater migration and faster functional maturation when treated with vascular factors, SPARC and SerpinE1, suggesting that the maturation of CNS vasculature may similarly regulate the timing of human fetal interneurons migration. In support of this, the MGE (also known as the germinal matrix) is the primary site periventricular hemorrhages (PVH) in prematurely-born babies (Luo et al., 2019). It has been hypothesized that this is due to newly formed (and thus weaker) vasculature in the MGE during the third trimester (Ballabh et al., 2004; Ballabh et al., 2007). As such, it is an appealing possibility that the timing of vascularization helps to coordinate species-specific aspects of neuronal maturation. Here, delayed vascularization in humans may serve a regulatory role in delaying interneuron migration until pyramidal neurons have achieved sufficient maturity to allow for interneuron circuit integration.

We identified SPARC and SerpinE1 as important proteins that account for most of the EndoCM activity. Both factors are known to be secreted by ECs, and SPARC has been implicated in cell differentiation (Hrabchak et al., 2008; Stary et al., 2005) and migration such as regulation of the epithelial-to-mesenchymal transition and cancer metastasis (Arnold and Brekken, 2009). SerpinE1 (also known as plasminogen activator inhibitor-1, PAI-1) is also implicated in cell migration through the uPA/Urokinase pathway (Mahmood et al., 2018), which has been shown to control interneuron tangential migration by acting through the c-Met receptor and hepatocyte growth factor(HGF) (Powell et al., 2001). This raises the possibility that SerpinE1 effects may intersect with the HGF/c-Met pathway. Future studies will elucidate the pathways activated by SPARC and SerpinE1 to promote interneuron migration and functional maturation.

Of note, we empirically determined that human ventral telencephalic organoids require approximately 14 days pre-treatment with SPARC and SerpinE1 before we could detect enhanced migration *in vitro*. This may reflect a human-specific difference in responding to endothelial cues compared with mouse interneurons that operate under a more compressed timescale. In fact, this may account for previous observations by Nicholas and colleagues that migration of hSC-interneurons into host mouse cortex occurs at a slow rate over 7 months (Nicholas et al., 2013). This may be explained by host vascularization of the xenograft, which has been shown to occur over a period of months (Mansour et al., 2018). We hypothesize that pretreatment with SPARC and SerpinE1 allows our xenograft to bypass host vascularization as a means to trigger migration. Therefore, pretreatment may prime hSC-interneurons to more rapidly integrate and mature into the host cortex.

Our findings demonstrate the utility of priming hSC-interneurons for transplantation by pre-inducing them to adopt a migratory state. Although, there are likely other impediments to overcome before hSC-interneurons are fully synchronized with the developmental timescale of host mouse cortex, our findings represent an important step towards harnessing the full potential of hSC-interneurons. Cortical interneurons are critical modulators of brain function (Kepecs and Fishell, 2014; Tremblay et al., 2016) and it is important to develop human interneuron models of disease (Catterall, 2018; Inan et al., 2013; Marin, 2012; Rapanelli et al., 2017). Moreover, our pre-treated hSC-interneurons also exhibit more maturity *in vitro*, suggesting SPARC/SerpinE1 treatment may also be effective when applied to organoid and monolayer approaches. Finally, consistent with previous work (Karakatsani et al., 2019), paracrine cues derived from ECs may similarly regulate the development of other neuronal populations. As such, our study may well serve as a general strategy for inducing hSC-neuron functional maturation in other neuronal cell types such as pyramidal neurons.

---

**Supplemental Figure 1.**
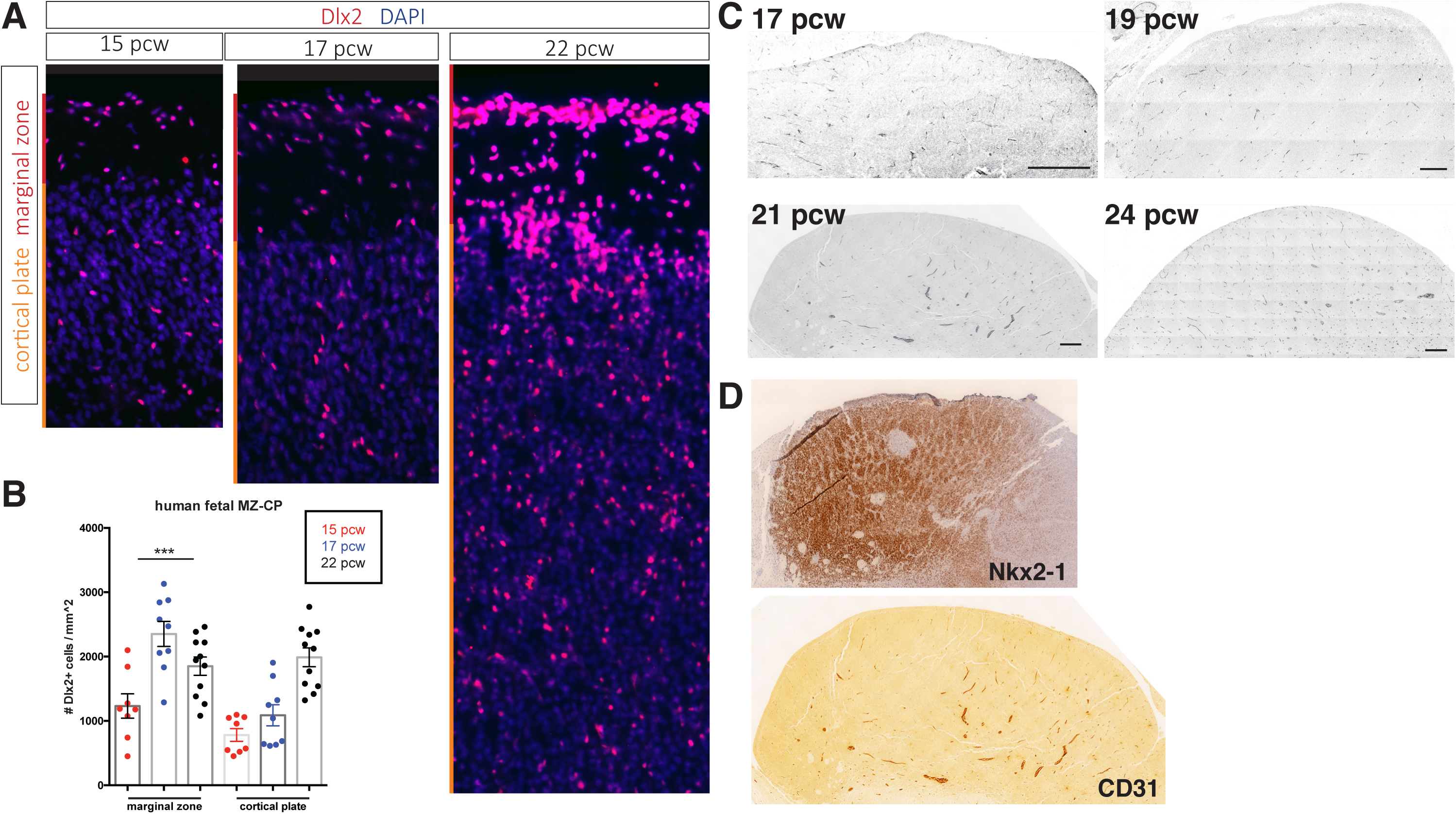
Cortical interneurons populate the human fetal cortex over a protracted period. (A) Representative immunostaining for Dlx2 (red) in coronal sections of human fetal brain at 15, 17 and 22 post-conception weeks (pcw). Red bar denotes marginal zone (MZ), orange bar denotes cortical plate (CP). (B) Quantification of Dlx2+ interneuron cell density/mm^2^. (c) Dlx2 cell density (/mm^2^) present in MZ and CP. (C) Transverse sections of human fetal MGE at 17, 19, 21 and 24 pcw immunolabeled for CD31. Scalebar denotes 200 microns. (D) Adjacent sections of 21 pcw MGE shows Nkx2-1 expression.

**Supplemental Figure 2.**
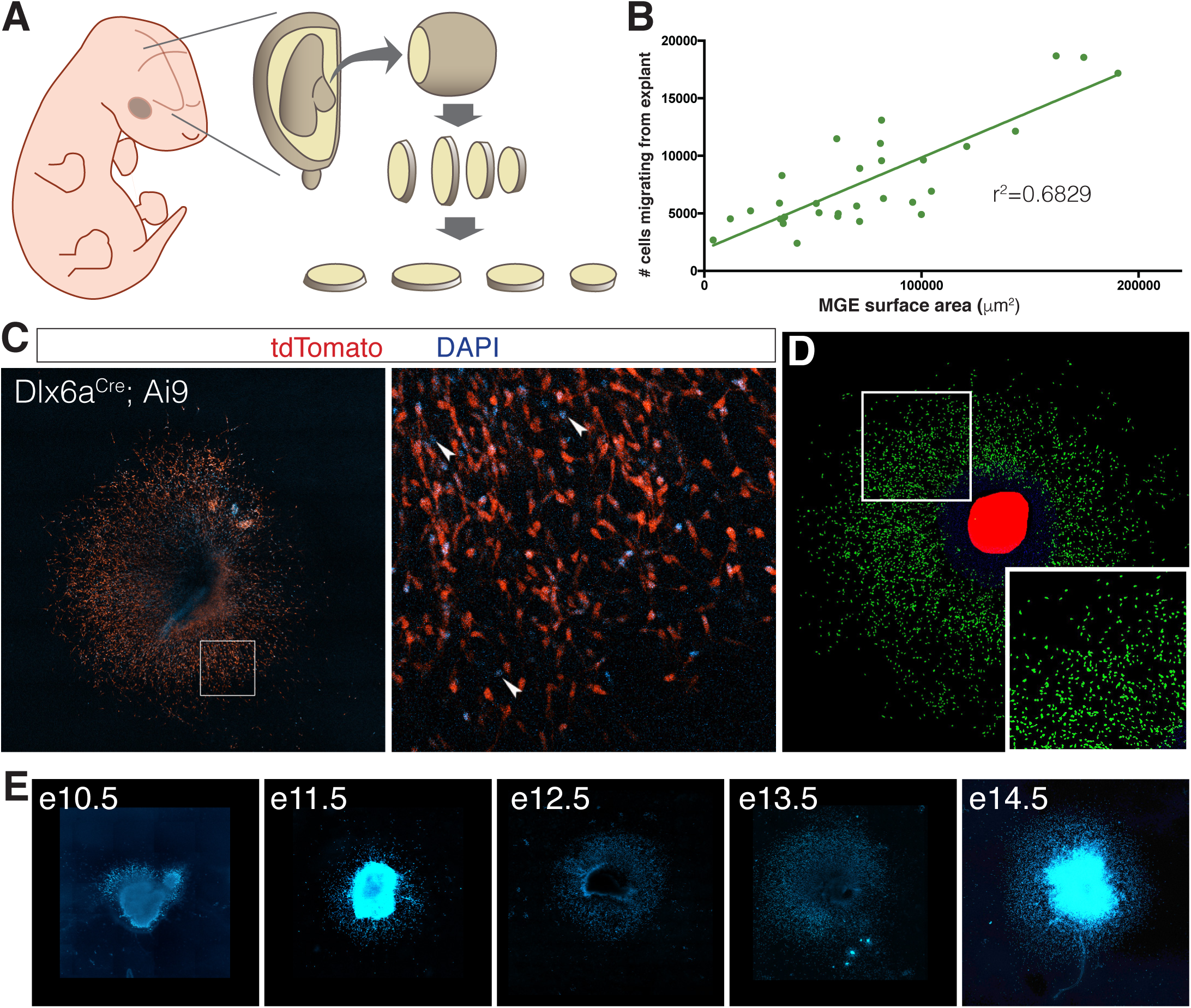
MGE explants cultured from mouse embryos at various ages. (A) Schematic representation of MGE explant preparation. MGE is dissected from embryo and sectioned at 250 microns. (C) Interneuron migration in e11.5 MGE explants linear correlation with surface area of MGE explant. (C) e14.5 MGE explant cultured from Dlx6a^Cre^; Ai9 embryo. Right, higher magnification of boxed region on left. Arrowheads show DAPI+ cells that are tdTomato-negative. (E) Representative DAPI-labeled MGE explants from embryos ages e10.5 to 14.5.

**Supplemental Figure 3.**
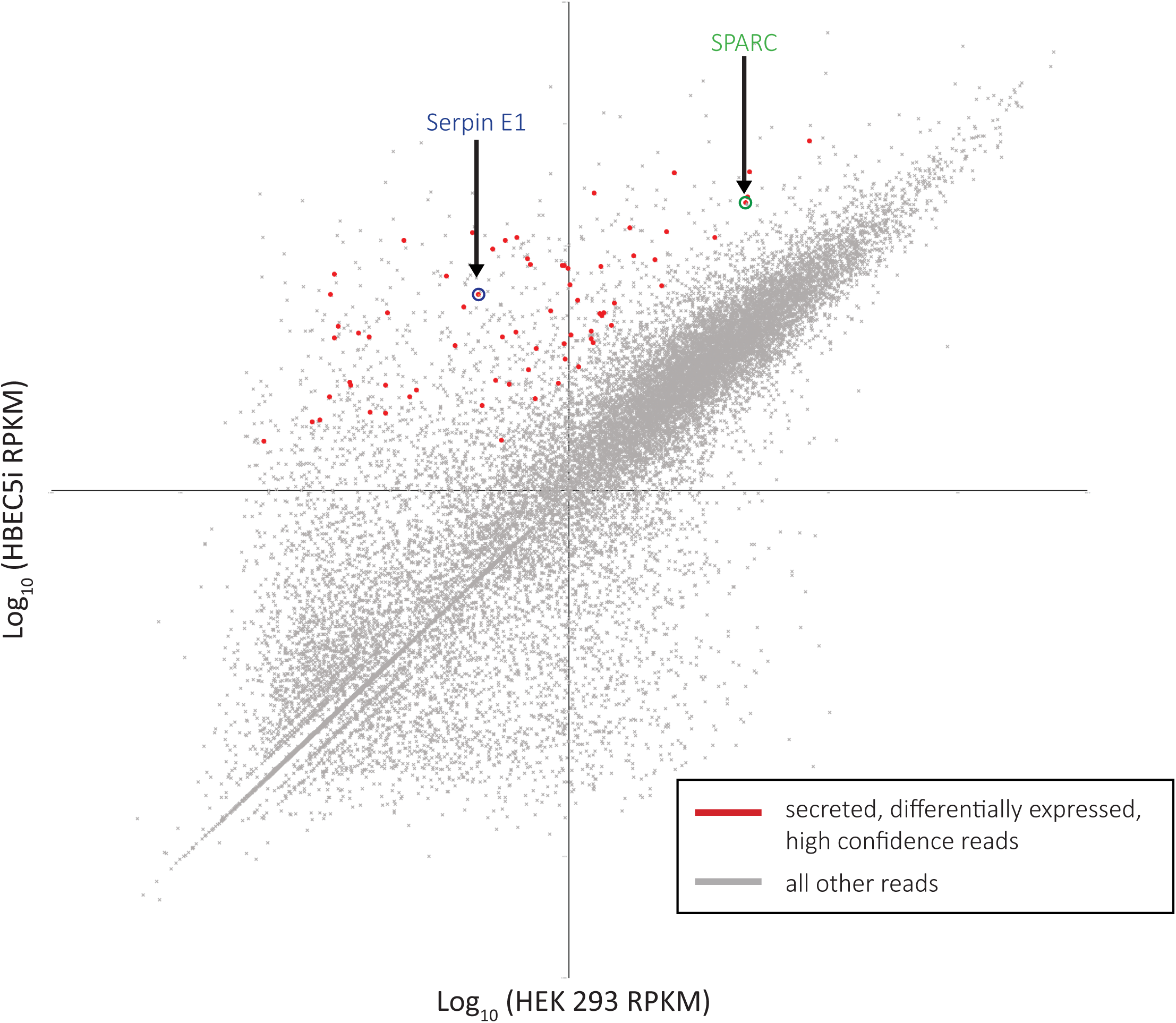
RNA-sequencing analysis identifies SPARC and SerpinE1 as highly enriched secreted factors in EndoCM versus HEK CM. Scatterplot of RNA-seq reads plotting Log_−10_(HEK 293) sample versus Log_10_(HBEC5i) sample. Red dots are HBEC5i-enriched secreted factor hits. All other reads are grey dots. Serpin E1 circled in blue. SPARC circled in green.

**Supplemental Figure 4.**
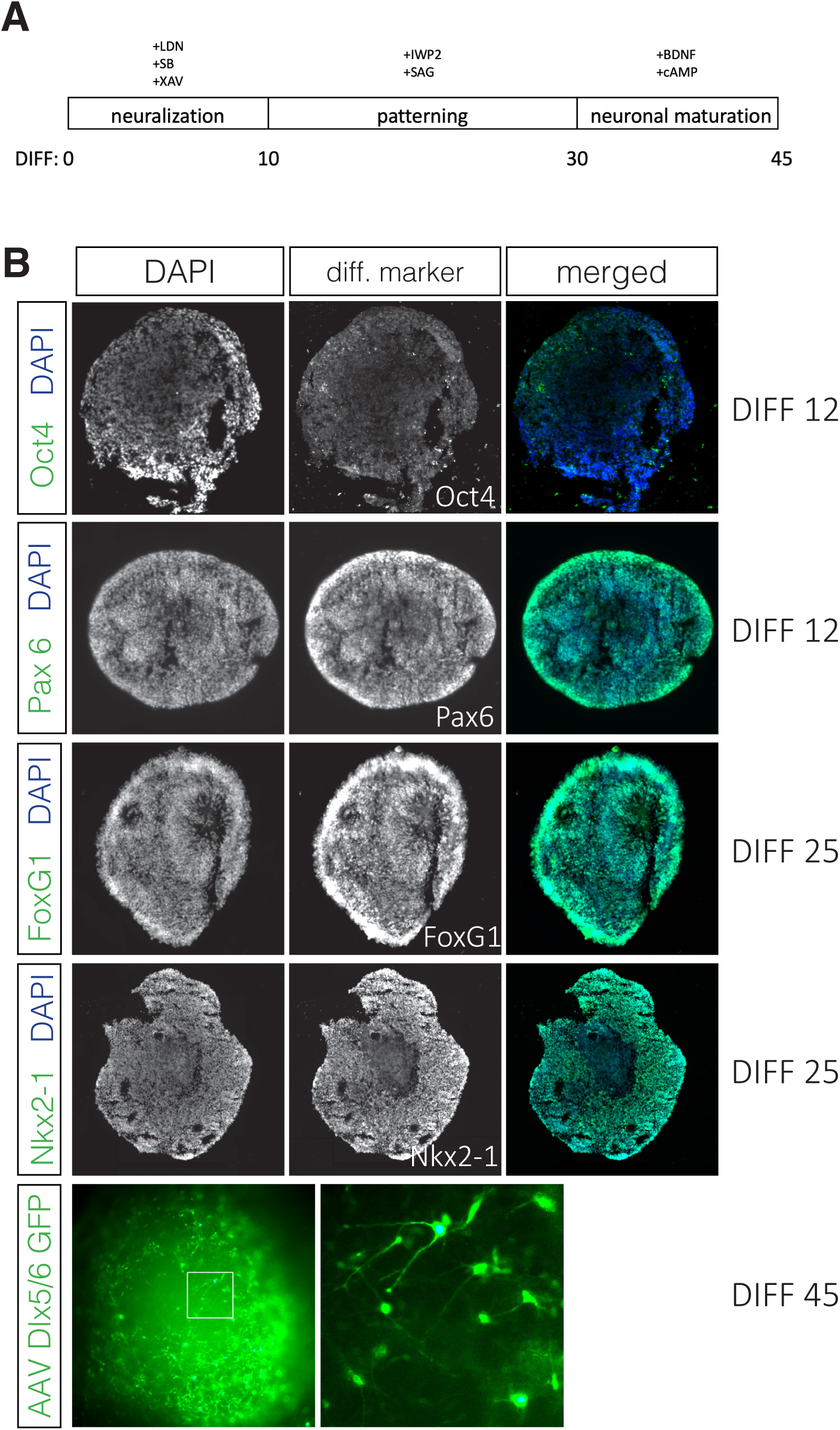
Human stem cells efficiently differentiate into interneurons. (A) Time line for hSC differentiation to cortical interneuron fate. (B) Differentiation day (DIFF) 12 organoids sectioned at 16 microns are largely Oct4-negative and almost completely Pax6-positive, indicating successful neural differentiation. DIFF 25 organoids sectioned at 16 microns are FoxG1- and Nkx2-1-positive. DIFF 45 whole organoid labeled with AAV-Dlx5/6-eGFP shows efficient differentiation to interneuron cell fate.

**Supplemental Figure 5.**
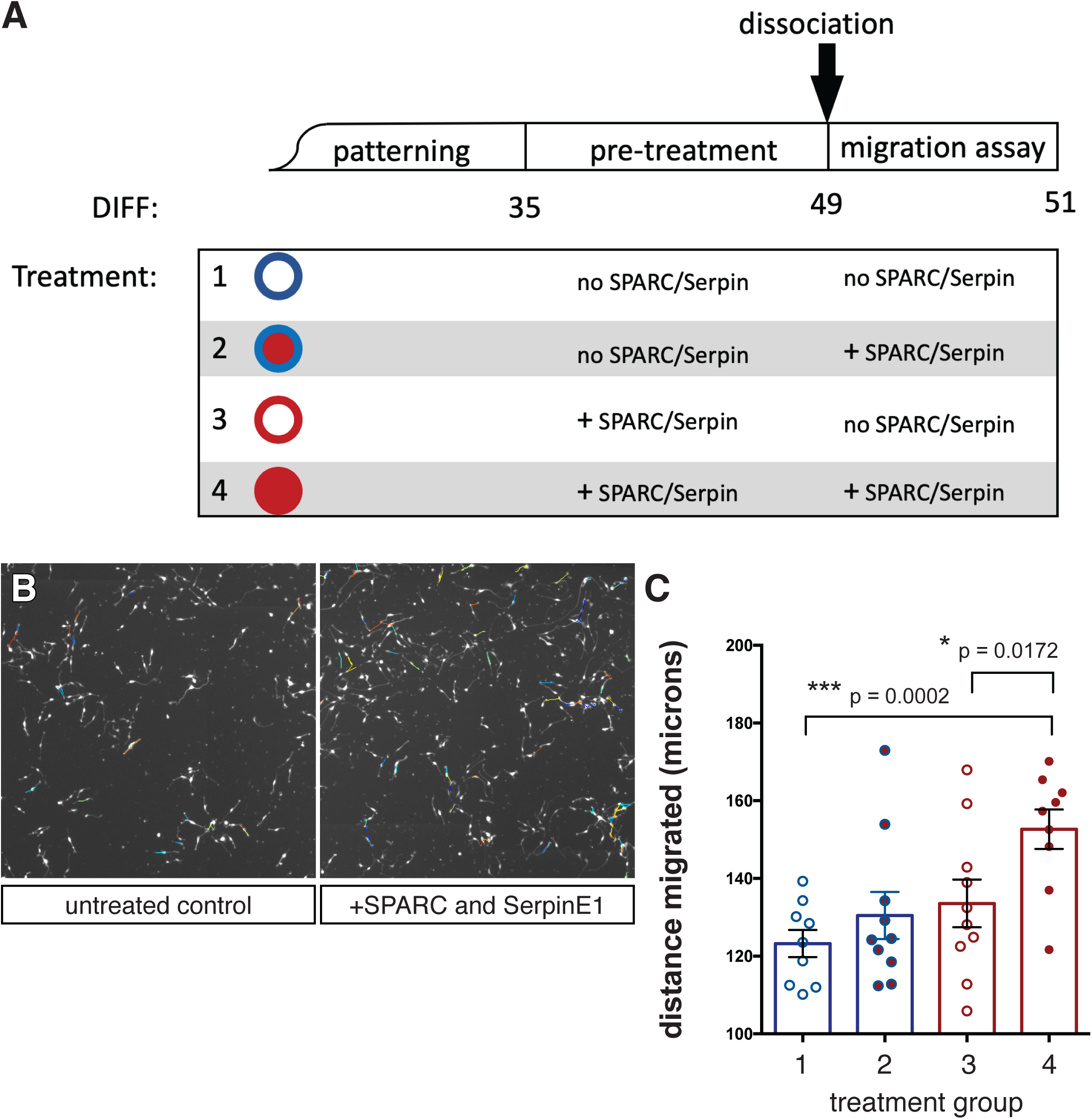
Human stem cell-derived interneurons migrate further in vitro with SPARC and SerpinE1 treatment. (A) Schematic of *in vitro* treatment groups for hSC-interneurons with SPARC and SerpinE1. (B) Example image of live tracking of hSC-interneurons plated as monolayer. (C) Quantification of hSC-interneuron treatment groups for total migratory distance over 48 hours. Paired t-test, * p < 0.05; **p < 0.01; *** p < 0.001

**Supplemental Figure 6.**
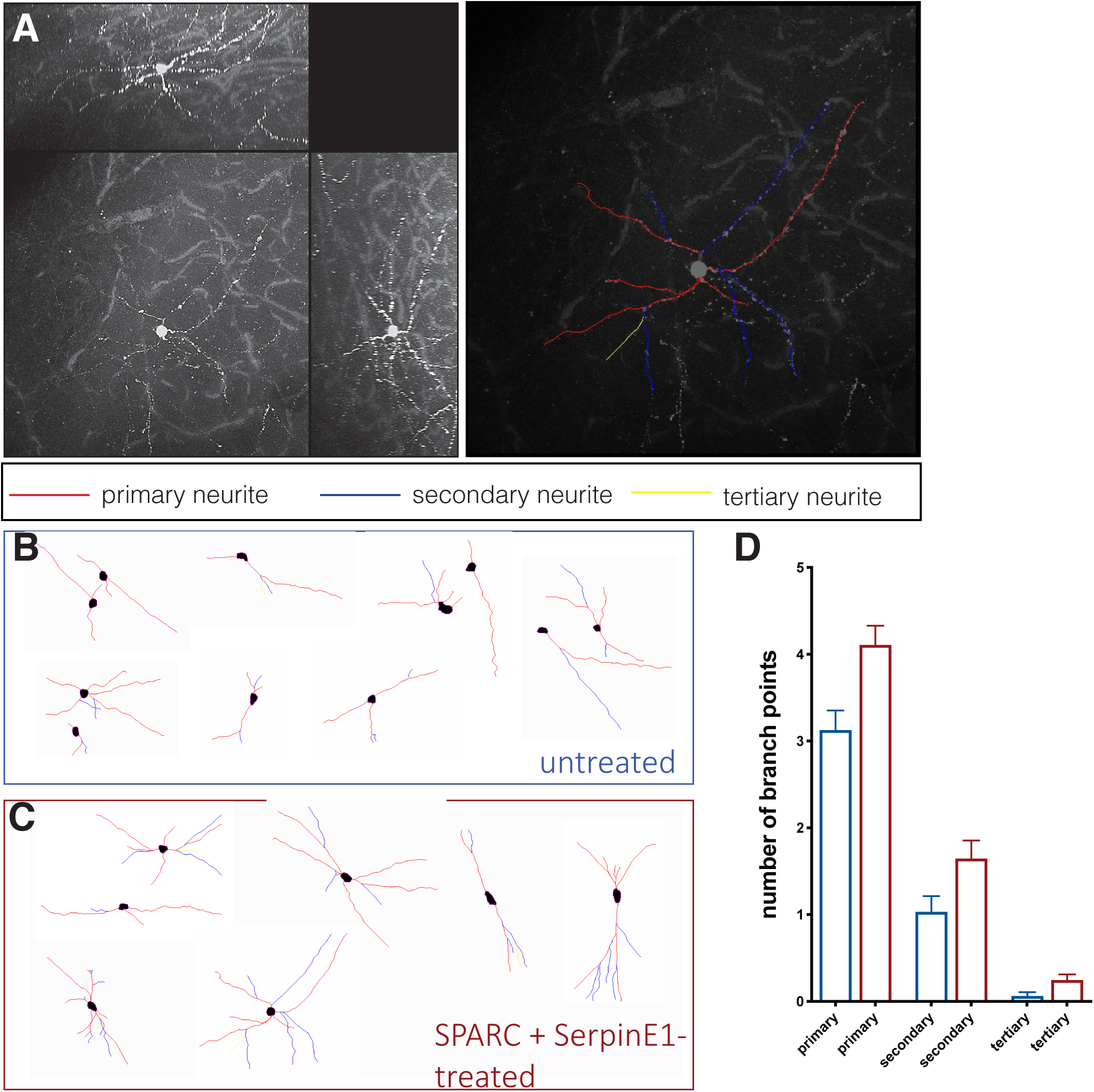
SPARC and SerpinE1-pretreated hSC-interneurons exhibit more complex morphologies after xenotransplantation into host mouse cortex. (A) Example of traced hSC-interneuron in cortex 2-months post-transplant. Primary neurite in red, secondary neurite in blue, tertiary neurite in yellow. (B, C) Representative traces of cell morphology for (B) untreated control and (C) SPARC/SerpinE1 pre-treated hSC-interneurons 2-months post-transplant. (D) Quantification of branch points in primary, secondary and tertiary process for untreated control and SPARC/SerpinE1 pre-treated groups.

**Supplemental Figure 7.**
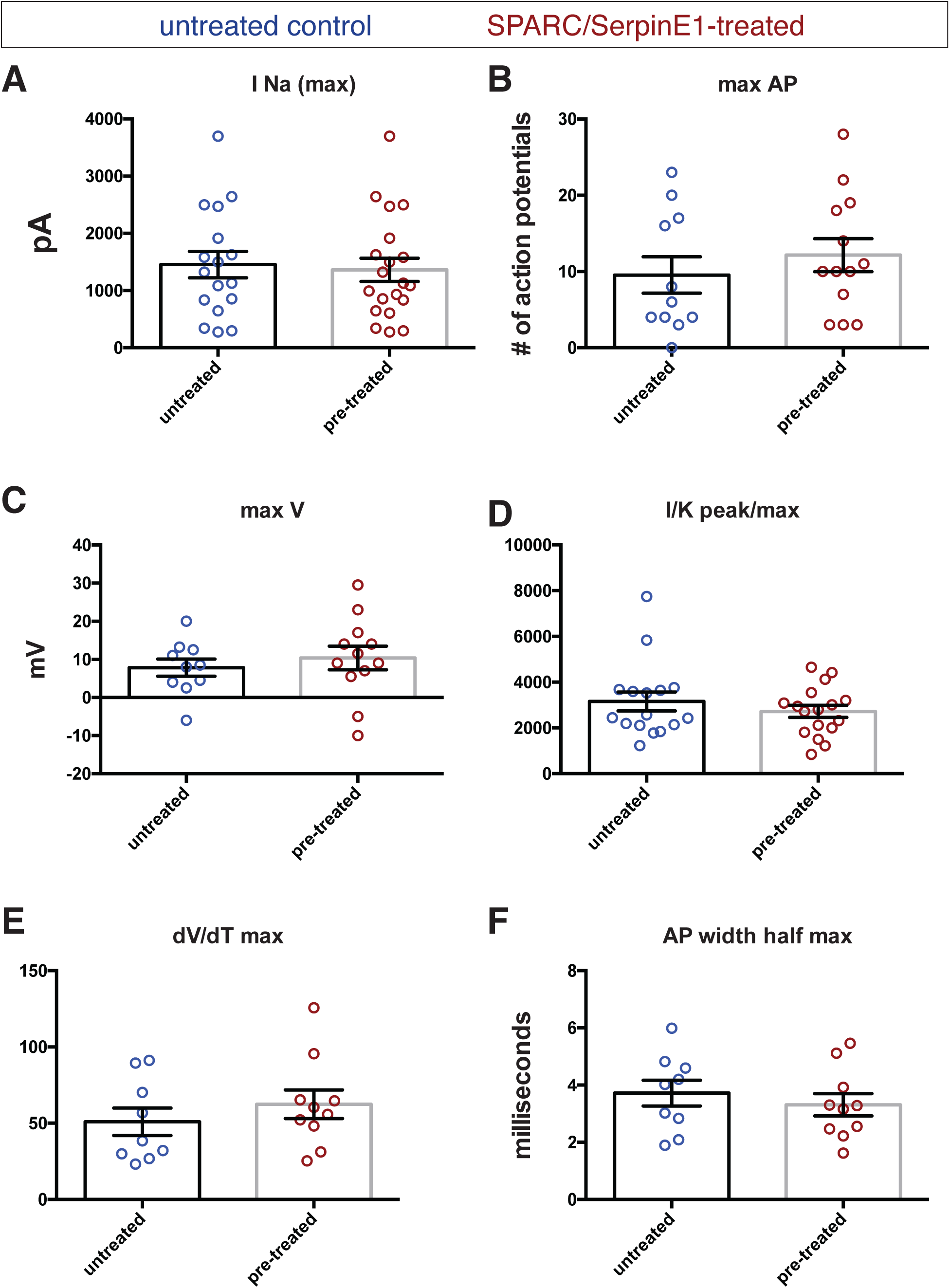
Electrophysiological recordings of control and pre-treated hSC-interneurons shows pre-treated group trends towards greater maturity by most measures. Untreated control (blue) and pre-treated (red) hSC-interneurons measured for (A) Sodium current (pico-Amps), (B) maximum number of action potentials (APs) elicited by square pulse depolarization, (C) maximum AP depolarization (milli-Volts), (D) Sodium-to-potassium peak current/max AP depolarization, (E) AP risetime, (F) AP width at half maximum amplitude (milliseconds).

## Materials and Methods

### MGE Explant Migration Assay

Swiss Webster timed plugs were generated to obtain embryos ranging in age from e10.5 to e15.5. MGE was microdissected under sterile conditions into iced cold Leibovitz’s L15 medium (Gibco). MGE tissue was then quickly embedded into low melt agarose (4 %) in Leibovitz’s L15 medium (Gibco). Embedded blocks were sectioned at 250 microns using a vibratome (Leica VT1000). MGE slices were then individually transferred into 4-well slides then covered with 300 uL Matrigel (Corning) (1:1 dilution) in Neurobasal medium (Gibco). Slides were then transferred to 37°C incubator for 15 minutes to allow Matrigel to solidify. Then, 300 uL MGE medium containing (Neurobasal medium, 2% B27, 1% N2, 1% Glutamax and 1% Penicillin/Streptomycin) was added on top of Matrigel. After 48 hours, MGE explants were fixed at room temperature in 4% paraformaldehyde/PBS for overnight and DAPI (2 ng/mL) was added to the chamber to label explants. Following 1 washed with PBS, MGE explants were imaged by confocal microscopy. Confocal images were analyzed using Image J plugin, a combination of background removing, cell counting and maximum intensity in order to automatically segment and quantify DAPI+ nuclei.

The following reagents were added to MGE explants diluted in overlying Matrigel:

| Reagent | Working concentration |
| --- | --- |
| p-EndoCM | 1:1 |
| EndoCM | 1:1 |
| SPARC | 50 ng/mL |
| SerpinE1 | 20 ng/mL |
| SPARC function-blocking antibody (mab 303 (Sweetwyne et al., 2004)) | 1:5 |
| SK216 (Masuda et al., 2013) | 100 uM |

For untreated controls, the appropriate vehicle at the same volume was added to x medium. Vehicle for p-EndoCM and EndoCM: Neurobasal Vehicle for SPARC: water Vehicle for SerpinE1: PBS Vehicle for Sparc function-blocking antibody: PBS; Vehicle for SK216: water.

### Organotypic Slice Culture Migration Assay

We employed a modified protocol previously described (Baffet et al., 2016). Briefly, Dlx6a^Cre^; Ai9 e11.5 embryos were sac’d and whole brain was isolated by microdissection in aCSF medium bubbled with O_2_. Brains were embedded in low melt agarose (4 %) in aCSF medium and sectioned by vibratome at a thickness of 250 microns. Sections were imaged live daily by confocal microscopy. Images were analyzed using Image J plugin trackMate in order to automatically segment and quantify migrating interneurons in cortex.

### Derivation of Conditioned Medium

Primary culture endothelial cells were derived as previously described (Baffet et al., 2016) and maintained in complete mouse endothelial cell medium from Cell Biologics. To collect p-EndoCM, cells were rinsed with PBS and then medium was switched to Neurobasal and 1% Glutamax for 3 days. Supernatant was collected, centrifuged at 4,000 × g for 15 minutes and filtered to remove cells. Then medium was concentrated using a 3000 kDa MWCO Centricon column (Millipore). Concentrated medium was stored at −80°C until use. Similarly, HBEC-5i cells were maintained in (DMEM/F12, 10%FBS, 40 ug/mL Endothelial Cell Growth Supplement) medium, rinsed with PBS and then switched to Neurobasal, 1% Glutamax medium for 3 days. EndoCM was similarly spun down, filtrated and concentrated using 3000 kDa MWCO Centricon column (Millipore). Concentrated EndoCM was also stored at −80°C until use. For size fractionation experiments, 10, 50 and 100 kDa MWCO Centricon columns (Millipore) were used for media concentration. HEK 293 cells were maintained in DMEM, 10%FBS, 1% Glutamax medium and switched to Neurobasal, 1% Glutamax medium following PBS rinses to collect HEK CM. It was also concentrated using 3000 kDa MWCO Centricon columns.

### RNA Sequence Analysis

HBEC-5i and HEK 293 cells were switched over to Neurobasal, 1% Glutamax medium in the same manner as if EndoCM and HEK CM were to be collected. 3 days later, cells were harvested in Trizol and RNA was purified using directZol miniPrep kit (Zymo). Samples were quality controlled and sequenced at Novogene using Illumina HigSeq/MiSeq with a sequencing depth of 20 million reads. Sequence data was then analyzed using DAVID to identify high confidence differentially-expressed hits of the appropriate molecular weight. These data were further screened by Gene Ontology search term “Extracellular Space” to identify secreted factors. Additional curation was performed to remove membrane-tethered proteins and pseudogenes in order to arrive at a short list of candidates (Table 1).

### Immunohistochemistry and Western Blots

Immunohistochemistry on *Apcdd1* mutants were as described previously(McKenzie et al., 2019). E14.5 Apcdd1 mutant and wildtype control brains were cryosectioned at 16 microns and immunolabeled with Alexa488-tagged Isolectin (1:100; Thermo Fisher Scientific); Calbindin (1:1,1000; ImmunoStar). Sections were acquired as tiled maximum projection images for analysis using FIJI for blood vessel quantification and Image J plugin cell counting for segmentation and calbindin+ cell counts. Western blot antibodies: Goat-anti-SPARC (1:1000; R&D Systems) and Rabbit-anti-SerpinE1 (1:1,000; Abcam) were used to detect EndoCM and HEK CM concentrated medium, both loaded with 20 uL. Xenograft 50 micron sections were immunolabeled with anti-Dlx2 (1:1000; Millipore).

### Human Fetal Cortical Tissue

We obtained fetal tissue samples for research following induced termination of pregnancy for maternal indications. Sample collection followed the policies of the Columbia University Irving Medical Center Institutional Review Board. IRB waiver AAAS5541 was obtained for non-human subjects research, deemed medical waste.

### Human Stem Cell Differentiation

Parental human iPSC line (NCRM 1, NIH Common Fund Regenerative Medicine Program) was genetically altered by CRISPR-mediated targeting of AAVS1 locus to introduce floxed-stop CAG-boosted tdTomato donor DNA construct. EF1-alpha promotor-driven Cre recombinase was introduced episomally to generate pan-tdTomato+ human iPSC line (pan-red line). Pan-red line was differentiated towards ventral telencephalic fate using and adaptation of previously described methods (Bagley et al., 2017; Xiang et al., 2017). In brief, pan-red hiPSCs were plated into ultra-low attachment u-bottom 96-well plates (9,000 cells/well) in neural induction medium containing LDN-193189 (100 mM), SB431542 (10mM) and XAV939 (10mM) to form organoids for 10 days. From day 10-17, cells changed to neuronal differentiation medium containing N2 and B27 (Invitrogen) with IWP2 (2.5 uM) and SAG (100nM). From day 18 onwards, neuronal differentiation medium also contains BDNF (20 ug/ml) and cAMP (125 mM) and ascorbic acid (200 ug/ml). At day 35, SPARC (50 ng/mL) and SerpinE1 (20 ng/mL) is added for 14 days. At day 49, organoids are gently dissociated in EDTA for 5 minutes, then Accutase 15min at 37C for downstream experiments (*in vitro* migration assays and xenotransplants).

### Xenograft Migration and Morphology Analysis

We performed stereotactic intracranial injections into the right frontal subcortical white matter (coordinates from Bregma: 1.0 mm right – 1.0 mm front – 1.0 mm deep) of NRG (NOD.*Cg-Rag1^tm1Mom^Il2rg^tm1Wjl^*/SzJ, Jackson Laboratories) mouse pups at p6-9 as previously described, adapted for mouse pups(Lei et al., 2011). We injected 12 P5/P6 NRG pups with control and 12 P5/P6 NRG pups with SPARC/SerpinE1-treated cells (∼50 000 cells/injection). At 28- and 56-days post injection animals were sacrificed and brains were processed for analysis (below). All procedures were performed according to Columbia University IACUC protocol no. AC-AAAV0463.

1-month post-transplant, host cortex was sectioned by vibratome at 50 microns. Migratory distance was assessed by scoring tdTomato+ cell linear distance from graft site. 2-months post-transplant, 300 micron vibratome sections were imaged by confocal microscopy to visualize tdTomato+ cell morphology. Processes were traced using Image J Plugin NeuronJ to assess neurite length and degree of branching.

### Acute Brain Slice Preparation and Electrophysiology

A more complete description of acute brain slice preparation and basic electrophysiology methods has been previously described (Crabtree et al., 2016; Crabtree et al., 2017).

Briefly, mice were anesthetized with isoflurane, decapitated, and brains were removed quickly and chilled in ice-cold dissection solution, which contained the following (in mM): 195 sucrose, 10 NaCl, 2 NaH_2_PO_4_, 5 KCl, 10 glucose, 25 NaHCO_3_, 4 Mg_2_SO_4_, and 0.5 CaCl_2_, and was bubbled with 95%O_2_/5%CO_2_. Coronal brain slices (∼300μm) centered on the injection site of the transplanted stem-cell-derived neurons were cut using a vibratome (a region covering the PFC through the region ∼1mm caudal to the corpus callosum). After slicing, brain slices were immediately transferred to a recovery chamber and incubated at room temperature in recording solution for a minimum of 30 minutes before recording. Total time between decapitation and procedure end was typically 12-16 minutes.

At the time of recording, slices were transferred to a submerged recording chamber and continuously perfused with standard aCSF (Crabtree et al., 2016). Whole-cell patch-clamp recordings were made using borosilicate glass pipettes (initial resistance, 2.0–5.5 MΩ). An internal solution was used that contained the following (in mM): KMeSO_4_ 145, HEPES 10, NaCl 10, CaCl_2_ 1, MgCl_2_ 1, EGTA 10, Mg-ATP 5, and Na_2_GTP 0.5, pH 7.2 with KOH. Solution junction potentials were small and were not corrected.

#### Basic electrophysiology

Recordings employed an Axon 700B MultiClamp amplifier, CV-7B headstage, and a Digidata 1440A data acquisition system. All signals were acquired at 10kHz (100µs). With the exception of spontaneous synaptic recordings (filtered at 2kHz), all other signals were filtered at 10 kHz. Cells targeted for recording were identified by red fluorescence. Confirmation of correctly targeted recordings was further validated by observation of a significant reduction in red fluorescent signal in the recorded cell at the end of the recording likely resulting from “wash-out” of the indicator protein via the recording pipette solution.

#### Current clamp recordings

All cells were forced to −70mV with a small negative current of variable amplitude. Bridge-balance mode was employed to minimize voltage errors and artifacts.

*Resting membrane potential* was reported as the cell voltage in I=0 mode observed shortly after whole-cell membrane rupture. The majority of cells were silent at rest leading to an uncomplicated reporting of V_rest_. A minority of cells (typically more hyperpolarized cells), however, displayed spontaneous AP or AP-like events rendering V_rest_ measures somewhat ambiguous. In this subset, V_rest_ was estimated as the midpoint voltage between the AP threshold voltage and deepest hyperpolarization after the AP.

#### Membrane time constant

Using small hyperpolarizing current steps, the region of the voltage response from 5ms after the start of the step to 205ms within the step was fitted with a single exponential using the standard Clampfit Chebyshev-method fitting routine. Accuracy of fits were further confirmed visually.

*Action potentials* were elicited in cells forced to −70mV with small (2.5-5pA), incremental current steps of 500ms duration. Unless indicated otherwise, reported metrics assessed the first AP elicited from current step recordings.

#### Action potential width

The first AP elicited from current step recordings was used for analysis. Action potential widths were measured using the standard Clampfit analysis routine “half-width,” the action potential width at half-height, and reported as AP width.

#### Action potential threshold

The first AP elicited from current step recordings was used for analysis. The voltage trace of this current step was converted to a time versus dV/dt plot and overlaid onto the original AP voltage trace. AP threshold was then determined visually as the first significant deviation of dV/dt from its baseline rate. All traces analyzed were assessed together in a single analysis session and all traces were displayed at the same time and dV/dt scale to avoid bias in threshold detection. A subset of traces were converted to “phase-plane” plots (V versus dV/dt) to further validate threshold assignments.

*Action potential maximum rate of rise* (dV/dt max) was determined using the standard Clampfit “maximum rise slope” routine. As with other AP measures, the first AP elicited from current step recordings was used for analysis.

#### Voltage clamp recordings

All cells were held at −70mV unless otherwise noted. As cell resistances were high (typically ∼1-2GΩ), pipette series resistances were low (typically < 10MΩ), and maximal elicited currents were relatively small (typically 1-3nA), series resistance compensation was not employed. The cell capacitance reported is that reported by the 700B amplifier which is the fast component which represents contributions from the soma and proximal process compartments.

*Voltage activated currents* were elicited with incremental 100ms voltage steps (from −100mV to +50mV) of 100ms duration in cells held at −70mV. Membrane resistance was determined from the current response of the voltage step to −80mV (ΔV, −10mV). Transient inward sodium currents were reported as the maximal sodium current elicited (I-Na_max_), typically observed at the step to −20mV or −30mV. Outward potassium currents were reported as the maximal potassium current elicited (I-K_max_) at the step to +50mV. The I-Na/I-K ratio was then derived from these two values.

*Spontaneous synaptic transmission* was recorded from cells held at −70mV in the absence of TTX. A standard recording duration of 3 minutes was employed. The observed synaptic events are likely dominated by glutamatergic synaptic events as the reversal potential for GABA-A currents of our solution combination was ∼-60mV. The frequency of synaptic events was highly variable between cells with some cells “silent” (or nearly so) while other cells had synaptic event frequencies in excess of 20Hz. Due to the extreme variability in these synaptic event profiles, herein we present only exemplary traces of the synaptic activity we observed.

#### Statistics

Electrophysiological parameters were compared with pairwise t-tests between conditions. T-tests were 1-tailed with a directional hypothesis of “more maturity” of the metric in the treated group.

